# Microbial community composition explains wintertime greenhouse gas fluxes in an oroarctic tundra ecosystem

**DOI:** 10.1101/2025.05.28.656102

**Authors:** Viitamäki Sirja, Eronen-Rasimus Eeva, Virkkala Anna-Maria, Maija E. Marushchak, Biasi Christina, Majamäki Renata, Igor S. Pessi, Hultman Jenni

## Abstract

Microbial communities play a central role in regulating the greenhouse gas balance in soil ecosystems. Microorganisms are active in the cold, snow-covered Arctic tundra soils; however, their contribution to the greenhouse gas budget during the Arctic winter remains poorly understood. To investigate the functional activity of bacterial and archaeal communities in oroarctic tundra soils during late winter, metatranscriptomic samples were collected alongside greenhouse gas (GHG) flux measurements across key vegetation types, which represent pH and moisture gradients. The transcription of central carbohydrate metabolism genes, various stress-related genes, and high carbon dioxide (CO_2_) fluxes evidenced active microbial metabolism in winter. Vegetation type, soil C/N ratio, pH, and water content explained the functional activity and microbial community composition during winter. The transcription of functional marker genes for methane oxidation and denitrification, coupled with flux data, suggests that shrublands and meadows act as methane (CH_4_) sinks in winter, while all vegetation types function as small nitrous oxide (N_2_O) sources. Our results further demonstrate that the soil microbial community has a significant impact on wintertime GHG emissions in the oroarctic tundra, thereby enhancing the explanatory power of a statistical model beyond that of abiotic environmental variables alone. This represents a promising step toward developing microbial-mediated models, which are crucial for improving predictions of ecosystem responses to climate change.

## INTRODUCTION

Climate change is causing rapid and severe changes in the Arctic (IPCC 2023), with recent estimates indicating that this region is warming almost four times faster than the rest of the world (Rantanen *et al*. 2022). The Arctic tundra contains approximately 1,700 billion tons of carbon, roughly one-third of the global soil carbon stock (Tarnocai *et al*. 2009; Hugelius *et al*. 2014). As the climate warms and permafrost thaws, some of this carbon will be released as CO_2_ and CH_4_ (Schuur *et al*. 2022; Maes *et al*. 2024). Increased emissions, even when partially offset by the increased plant carbon uptake (See *et al*. 2024), induce a positive climate feedback loop that amplifies global warming. These large-scale changes are ultimately governed by small-scale processes such as microbial activity, which drives the decomposition of organic matter and the release of greenhouse gases (GHGs). Therefore, a thorough understanding of microbial communities and their responses to environmental changes in the Arctic tundra is urgently needed (Cavicchioli *et al*. 2019; Jansson and Hofmockel 2020).

Recent research on microbial communities in Arctic soils focuses on the high Arctic regions of the USA (Mackelprang *et al*. 2011; Coolen and Orsi 2015; Hultman *et al*. 2015; Waldrop *et al*. 2023), Canada (Steven *et al*. 2008; Yergeau *et al*. 2010; Varsadiya *et al*. 2021), Siberia (Kobabe, Wagner and Pfeiffer 2004; Liang *et al*. 2019), Sweden, (Mondav *et al*. 2017; Woodcroft *et al*. 2018) and Svalbard (Hansen *et al*. 2007; Schostag *et al*. 2015; Xue *et al*. 2019). In contrast, sparsely vegetated and dry Arctic-alpine regions are underrepresented (Metcalfe *et al*. 2018; Bourquin *et al*. 2022). Our previous study showed that vegetation drives summertime microbial community structure and activity in tundra heaths (Pessi *et al*. 2022; Viitamäki *et al*. 2022). Although the Arctic winter lasts eight to ten months, microbiological studies focus on the short growing season (Poppeliers *et al*. 2022). Nevertheless, studies have demonstrated microbial activity (Brooks, Williams and Schmidt 1996; Rivkina *et al*. 2000; Mikan, Schimel and Doyle 2002), increased microbial biomass (Brooks, Williams and Schmidt 1998; Isobe *et al*. 2018), decreased litter mass, and increased nutrient mineralization (Hobbie and Chapin 1996; Schimel, Bilbrough and Welker 2004) in sub-zero soils.

Studies show decreased plant carbon uptake due to increased respiration during non-growing seasons (See *et al*. 2024), but direct evidence regarding microbial roles is lacking. A few studies of arctic and alpine soils outside the growing season indicate strong seasonality in microbial communities (Männistö, Tiirola and Häggblom 2007; Björk *et al*. 2008; Schostag *et al*. 2015; Männistö *et al*. 2024). Similarly, while most GHG flux data from the northern high latitudes are from the warm season (Vogt *et al*. 2024), there is accumulating evidence that winter emissions significantly contribute to annual soil CO_2_ and CH_4_ budgets, with small but persistent emissions observed during winter (Zona *et al*. 2016; Treat, Bloom and Marushchak 2018; Pedron *et al*. 2022; Arndt *et al*. 2023). Winter N_2_O fluxes are less studied, but cold-season emissions are significant in some regions (Wagner-Riddle *et al*. 2017; Gao *et al*. 2020).

This study investigated winter soil microbial communities and their role in GHG fluxes across key vegetation types: shrublands, meadows, and fens in the oroarctic tundra. Oroarctic, the high-latitude mountainous tundra, is characterized by treeless ericoid-graminoid-dominated vegetation, mountainous terrain, and deep snow in the winter (Virtanen *et al*. 2016). Samples for metatranscriptomes and *in situ* CO_2_, CH_4_, and N_2_O fluxes were collected at maximum snow depth. We hypothesize that microbial processes related to GHG dynamics and organic matter decomposition continue during winter, with small CO_2_ and N_2_O emissions and CH_4_ emissions from wetlands observed. We also hypothesize that wintertime GHG fluxes can be linked to microbial CO_2_, CH_4_, and N_2_O metabolism and that microbial gene transcription provides additional explanatory power beyond that of abiotic environmental variables alone.

## METHODS

### Study area

The study area is in Kilpisjärvi (Gilbbesjávri in Northern Sámi), in Sápmi in northwestern Finland, within a valley between the fells Saana (Sána; 69°02’ N, 20°51’E; 1029 m a.s.l.) and Korkea-Jehkas (Jiehkkáš; 69°04’N, 20°79’E; 960 m a.s.l.)(Figure 1). This oroarctic tundra region is detailed in Virkkala *et al*., 2024. Samples were collected over four days in April 2021 from four vegetation types: evergreen shrubs (6 samples), deciduous shrubs (5), meadows (5), and fens (3), as classified by the Circumpolar Arctic Vegetation map (Walker et al., 2005). Shrub vegetation is described by Viitamäki *et al*. (2022). Meadows and seasonally dry fens are tundra wetlands dominated by graminoids, with fens also including a range of characteristic wetland species, such as sphagnum. Meadow plots had a mineral-rich mixture of decomposed organic soil and peat, while fen plots contained some mineral soil mixed with peat. Most plots had a shallow soil layer (< 50 cm) above bedrock.

**Figure 1.**
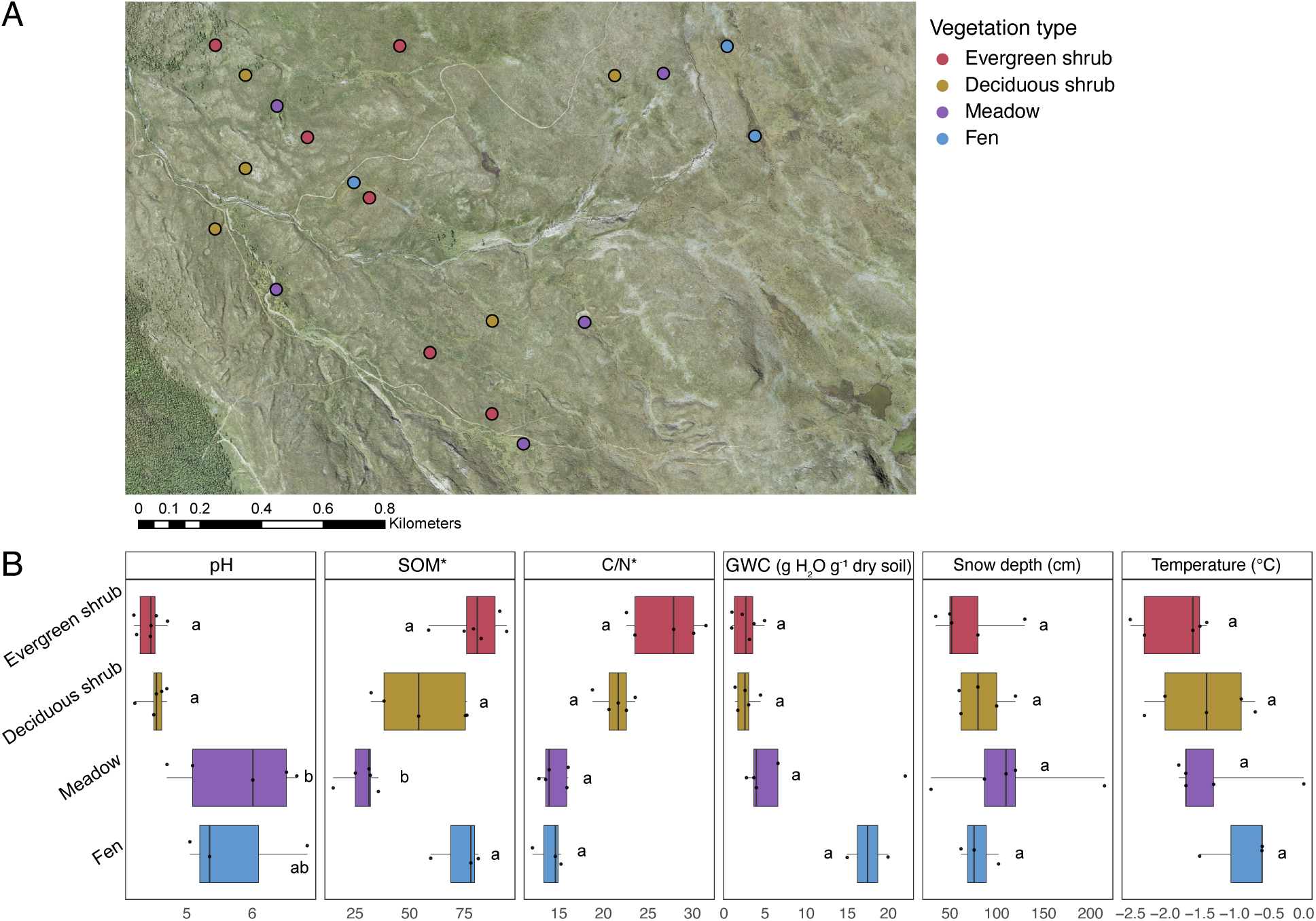
(A) Map showing the sampling plots by vegetation type. (B) Boxplot showing each vegetation type’s soil pH, SOM, C/N ratio, GWC, soil surface temperature, and snow depth. SOM and C/N ratio are measured in summer from the same plots (marked with *). Vegetation types followed by different letters are significantly different (Dunn’s test, p < 0.05). One unrealistically high outlier value for fen GWC was omitted.

### Data collection

#### Soil sampling for microbial data

Sampling plots were established in previous studies (Pessi *et al*. 2022; Viitamäki *et al*. 2022). Snow and aboveground plant parts were removed, and the topsoil was loosened with a sterile chisel and hammer. Samples were collected from the organic top layer beneath plant roots, targeting a depth of 5 cm. In fen plots, the hard, icy topsoil prevented reaching the target depth, so samples were taken from the deepest point possible, potentially increasing plant material content. Samples were collected with a sterile spoon, flash-frozen in dry ice, and stored at −80°C until nucleic acid extraction.

#### Supporting environmental data

Temperatures were measured from the snow pit just above the soil surface before sampling using a TENMARS TM-80N K/J Thermometer (Tenmars Electronics Co., Ltd., Taipei, Taiwan). Soil organic matter (SOM), carbon (C), and nitrogen (N) were measured from July 2018 soil samples (5-10 cm depth) from the same plots. pH and gravimetric water content (GWC) were measured from winter soil samples. Analyses followed Viitamäki et al. (2022), with a Leco 628 Elemental Analyzer (LECO Corporation, St. Joseph, MI, USA) used for C and N analysis. Two anomalously high GWC values (>20 g H₂O g⁻¹ dry soil) were replaced with the average vegetation type-specific GWC values, and one missing temperature value was replaced with the four-year mean April temperature 15 cm above the soil surface, measured with a TMS-4 microclimate logger (Niittynen et al., 2024).

#### Greenhouse gas flux data

Soil gas profiles and CO_2_, CH_4_, and N_2_O fluxes were sampled from the same plots as microbial samples using the snow gradient method (described in Marushchak *et al*. 2011). Gas samples of 25 mL were collected every 10 cm of the snowpack, injected into pre-evacuated 12 ml glass exetainers (Labco Exetainer, Labco Ltd.), and analyzed at the University of Eastern Finland, Kuopio, using a gas chromatograph using Agilent 7890B gas chromatograph (Agilent Technologies, Santa Clara, CA, USA), equipped with an autosampler (Gilson Inc., Middleton, WI, USA) and a thermal conductivity detector for CO_2_, a flame ionization detector for CH_4_, and an electron capture detector for N_2_O. Standard gas mixtures were analyzed with each batch to calculate gas concentrations.

### Data processing

#### Nucleic acid extraction and sequencing

RNA extraction followed a modified hexadecyltrimethylammonium bromide, phenol-chloroform, and bead-beating protocol (Griffiths et al., 2000; DeAngelis et al., 2009) using the AllPrep DNA/RNA Mini Kit (Qiagen, Hilden, Germany). Complementary DNA (cDNA) libraries were constructed using the NEBNext Ultra II Directional RNA Library Prep Kit for Illumina and NEBNext Multiplex Oligos for Illumina (New England Biolabs, Ipswich, MA, USA). Methods for extraction, as well as RNA and cDNA quality and quantity control, are described in Viitamäki et al. (2022). Paired-end sequencing was conducted on an Illumina NovaSeq S4 2×150 bp (Illumina, San Diego, CA, USA) at Novogene Co., Ltd., UK, yielding an average of 170M reads per sample.

#### Metatranscriptomic data processing and analysis

Sequence quality was assessed using FastQC v. 0.11.9 (https://www.bioinformatics.babraham.ac.uk/projects/fastqc) and MultiQC v. 1.12 (Ewels *et al*. 2016). Trimming and quality filtering were performed using Cutadapt v. 3.5 (Martin, 2011), with a quality cutoff of 20 and a minimum length of 50 bp.

SSU rRNA reads were removed using SortMeRNA v. 4.3.4 (Kopylova, Noé and Touzet 2012) to obtain protein-coding reads. The filtering utilized reference databases for bacterial and archaeal 16S and 23S, eukaryotic 18S and 28S, and 5S and 5.8S rRNA genes (https://github.com/sortmerna/sortmerna/tree/master/data/rRNA_databases). To analyze all protein-coding genes, filtered reads were mapped to the Kyoto Encyclopedia of Genes and Genomes (KEGG) Prokaryote database release 86 (Kanehisa and Goto, 2000) using DIAMOND v2.1.6.160 (Buchfink, Reuter, and Drost, 2021) with an E-value cutoff of 0.001. To analyze selected functional marker genes reads were mapped to a manually curated database (Leung and Greening 2020) using DIAMOND with an E-value cutoff of 0.001 and a query cover cutoff of 80%. Paired reads with a better E-value were chosen, and transcripts were normalized to transcripts per million (TPM) of rRNA-filtered reads.

For taxonomic analysis, trimmed reads were reduced to 10 million using seqtk v. 1.3 (https://github.com/lh3/seqtk) with a random seed 100. Active taxa were obtained using phyloFlash v3.4.2 (Gruber-Vodicka, Seah and Pruesse 2020) with the SILVA 138.1 SSURef NR99 database (main taxonomic ranks only), which was formatted for phyloFlash (https://zenodo.org/records/10047346). Taxonomic data were transformed into relative abundance and visualized using the R package Phyloseq (v. 1.46.0; McMurdie and Holmes 2013). Additional visualizations were conducted using R v. 4.3.3 (R Core Team 2020) and R packages from Tidyverse 2.0 (Wickham *et al*. 2019).

#### Flux calculation

To estimate winter GHG net flux between the snow and atmosphere, we applied Fick’s law of diffusion using measured CO_2_, CH_4_, and N_2_O concentration gradients within the snowpack. The gas flux calculation was based on the N_2_O concentration gradient between the lowest sampling depth in the snow and ambient air. The other concentrations were used as a quality control to ensure linear gradients. The diffusive flux was calculated as follows:

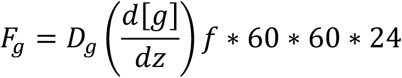

Where F_g_ is the gas flux (mg m^-2^ d^-1^), D_g_ is the gas-specific diffusion coefficient in snow (in cm^2^ h^-1^), dg/dz is the vertical concentration gradient (change in molar fraction per unit snow depth), M is the molar mass, and f is the snow porosity. The diffusion coefficient D_g_ was 0.139 cm^2^ s^-1^ for CO_2_ and N_2_O and 0.22 for CH_4_ (Sommerfeld, Mosier and Musselman 1993). Snow porosity was calculated by weighing a snow sample collected with a PVC tube (Ø 10 cm) and using the pure ice density (0.9168 g cm^−3^).

### Statistical analysis

#### Comparisons between vegetation types

We used the Kruskal-Wallis rank sum test and Dunn’s test with Bonferroni correction (R package FSA v0.9.5; Ogle *et al*. 2025) to compare environmental variables, microbial taxa relative abundances, and functional gene transcription between vegetation types. We performed Principal Coordinates Analysis (PCoA) to explore differences in microbial community composition and functional genes based on Bray-Curtis dissimilarity of taxonomic data (relative abundance) or functional data (TPM). Analyses were performed using *vegdist* (vegan package v2.6-6.1; Oksanen et al. 2025) and *cmdscale* functions. Data normality of gas fluxes was assessed with the Shapiro–Wilk test. A one-sample t-test was used for normally distributed data to determine whether fluxes differed significantly from zero. If the data were not normally distributed, the non-parametric Wilcoxon signed-rank test was applied instead. Unless otherwise specified, statistical tests were performed using the stats package (v4.4.3) in R (R Core Team, 2020).

To assess the influence of environmental variables on microbial community composition and functional genes, we conducted distance-based redundancy analysis (dbRDA) using the vegan package. The Bray-Curtis dissimilarity matrix was computed from the relative abundance or TPM data. dbRDA was performed with vegetation type, pH, SOM, C/N ratio, snow depth, and temperature as explanatory variables. Stepwise selection refined the model to include only significant predictors, resulting in vegetation type and pH. Variance inflation factor (VIF) confirmed minimal multicollinearity (VIF < 3).

#### Generalized additive models

We developed Generalized Additive Models (GAMs) to explore relationships between environmental factors, functional gene transcripts, and GHG fluxes. GAMs were selected to accommodate the nonlinear relationships between explanatory and response variables. Models were constructed using the mgcv package (v1.9.1; Wood 2025) with a Gaussian family and log link function to address the left-skewed distribution of TPM data (predominantly small values). Smooth term dimensions were constrained to three.

Explanatory variables included GWC, temperature, and C/N ratio, limited to these due to a small sample size (n=19). The C/N ratio correlated strongly with pH (r=-0.7, p=0.002) and SOM (r=0.56, p=0.001). Thus, C/N results also apply to pH (reversed) and SOM. We limited our analysis to genes *narG*, *nirK*, *norB*, [NiFe] hydrogenase type 1h, *nifH*, *coxL*, *pmoA,* and *mmoA* for their role in GHG and trace gas cycling. Note that most genes included in the GAMs are involved in CH_4_ and N_2_O production and consumption since CO_2_ is a central metabolite in many pathways and, thus, not specific to a single microbial gene or group. Plots lacking TPM data for specific genes were assigned zero values, assuming no transcription.

We developed two sets of models. First, we modeled each gene as TPM as a function of GWC, temperature, and C/N ratio to identify influential predictors of wintertime gene activity. Next, we modeled each GHG flux using the same environmental variables, then incorporated each gene’s TPM to assess how microbial gene data improve model performance and characterize gene-flux relationships.

Model performance was evaluated with adjusted R^2^ values from the GAM summary. Variable importance was quantified using a permutation approach in the vip package (Greenwell and Boehmke 2020), which assumes that randomly permuting important explanatory variable values in the training data would decrease model performance. The difference between baseline R^2^ and the R^2^ obtained after permutation served as our variable importance measure (higher values indicating greater importance). Relationship patterns (positive, negative, U-shaped, unimodal, or indeterminate) were characterized by examining response curves from the GAM output (see, e.g., Rissanen *et al*. 2021).

### Data availability

Sequences were deposited in NCBI SRA under BioProject PRJNA1265891 (SRA accessions SRR33650596–SRR33650614). The R code used for data processing and analysis in this study will be made available in the following GitHub repository upon publication: https://github.com/ArcticMicrobialEcology/kilpisjarvi-winter

## RESULTS

### Key soil properties — pH, SOM, and C/N ratio — differed significantly between vegetation types, while temperature and snow depth did not

Soil pH differed significantly between vegetation types (Kruskal-Wallis test, χ² = 12.572, df = 3, *p* < 0.01) (Figure 1B). Shrubs were significantly more acidic than meadows (Dunn’s test, *p* < 0.05). SOM varied between vegetation types (Kruskal-Wallis test, χ² = 12.938, df = 3, *p* < 0.01), with meadows having significantly lower SOM than evergreen shrubs and fens (Dunn’s test, *p* < 0.05). C/N ratio differed significantly between vegetation types (Kruskal-Wallis test, χ² = 14.905, df = 3, *p* < 0.01), with evergreen shrubs having significantly higher C/N than meadows and fens (Dunn’s test, p < 0.01). Water content was lower in shrubs than in meadows and fens, but without significant differences since the small sample size of fens (n = 2) limits the test’s statistical power. However, a p-value of 0.057 and a large effect size ε²=0.3 indicate a possible meaningful ecological difference. The soil surface temperature during sampling was −1.4±0.67°C (mean ± SD), and snow depth was 87.3±43.4 cm, without significant differences between vegetation types.

### All vegetation types were a source of CO2 and N2O, shrubs and meadows were methane sinks, whereas one fen plot was a CH4 source

CO_2_ emissions were detected in all soils, with deciduous shrubs exhibiting the highest values; however, no significant difference was found between vegetation types (Figure 2). Other vegetation types besides meadows had a flux significantly different from zero. In evergreen shrubs, the mean CO₂ flux was 607 µg CO_2_ m⁻² h⁻¹ (SD = 348, t(4) = 3.90, *p* = 0.0175), in deciduous shrubs it was 774 µg CO_2_ m⁻² h⁻¹ (SD = 334, t(4) = 5.18, *p* = 0.0066), and in fens it was 479 µg CO_2_ m⁻² h⁻¹ (SD = 166, t(3) = 5.78, *p* = 0.0103). A small uptake of CH_4_ was detected in shrubs and meadows, although the flux was not significantly different from zero. In fens, CH_4_ fluxes were close to zero, except for one plot exceeding 8 mg m^-2^ d^-1^. Generally, all vegetation types were minor sources of N_2_O, with a flux not significantly different from zero. CH_4_ concentrations typically decreased with depth in shrubs and meadows, indicating the consumption of atmospheric CH_4_ in the upper soil layers (Supplementary Figure S1). In contrast, such clear depth trends were not observed in fens, except for the high-emitting plot in winter.

**Figure 2.**
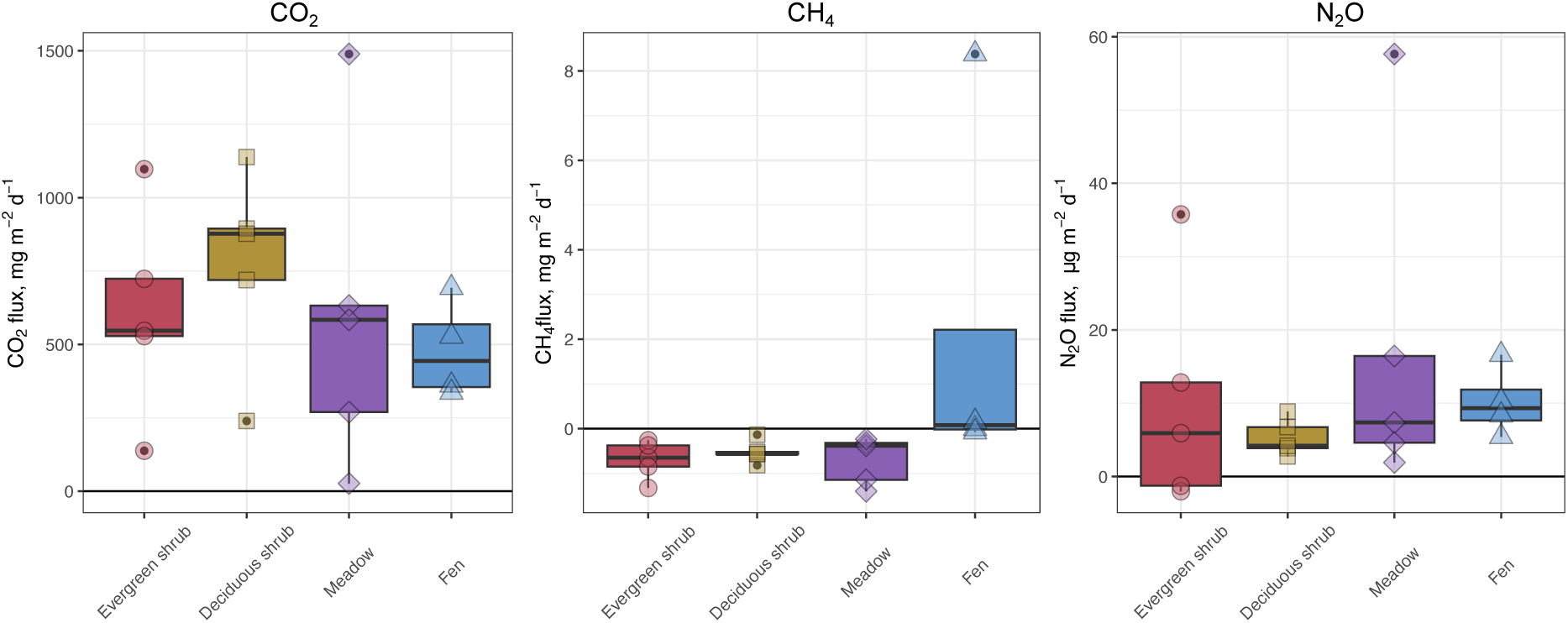
The GHG fluxes for CO2, CH4, and N2O were measured using the snow gradient method from the bottom of the snowpack to just above the soil surface for each vegetation type. Negative flux indicates gas consumption, and positive flux indicates emission to the atmosphere.

### Vegetation type and pH shape the active communities in winter

#### Bacterial community composition

Of the trimmed reads, 95% were rRNA. *Actinobacteriota*, *Alphaproteobacteria*, *Acidobacteriota,* and *Planctomycetota* represent approximately 75% of active taxa in shrubs, 65% in meadows, and 50% in fens, as cumulative relative abundance (Supplementary Figure 2). Community composition in shrubs was similar, whereas shrub communities differed from meadows and particularly from fens with higher pH and GWC. Most significant differences were found between evergreen shrubs and meadows, with *Acidobacterales*, *Bryobacterales*, *Vicinamibacterales*, and *Clostridiales* more active in evergreen shrubs (Kruskal-Wallis test, followed by Dunn’s test with Bonferroni correction; *p* < 0.05).

#### Bacterial community dynamics across vegetation types and environmental gradients

PCoA analysis showed that vegetation type explained variation in bacterial and archaeal community composition (Figure 3a) and functional genes (Figure 3b) in oroarctic tundra soils. In the dbRDA analysis for the taxonomic composition, the final model retained vegetation type and pH as predictors, explaining 65.78% of the total variance in community composition. The overall model was highly significant (F = 1.004, *p* < 0.001), and permutation tests showed that both pH (F = 12.197, *p* < 0.001) and vegetation (F = 4.907, *p* < 0.01) were significant contributors, with vegetation explaining 35.98% and pH explaining 29.81% of the variation. In the dbRDA analysis for the functional genes, the reduced model with vegetation type, pH, and temperature as predictors was highly significant (F = 2.801, *p* < 0.001) and explained 51.86% of the total variation. Vegetation explained 35.2% (F = 2.671, *p* < 0.01) and pH 9.3% (F = 2.617, *p* < 0.05) of the variation in the transcription of functional genes, whereas temperature did not have a significant contribution.

**Figure 3.**
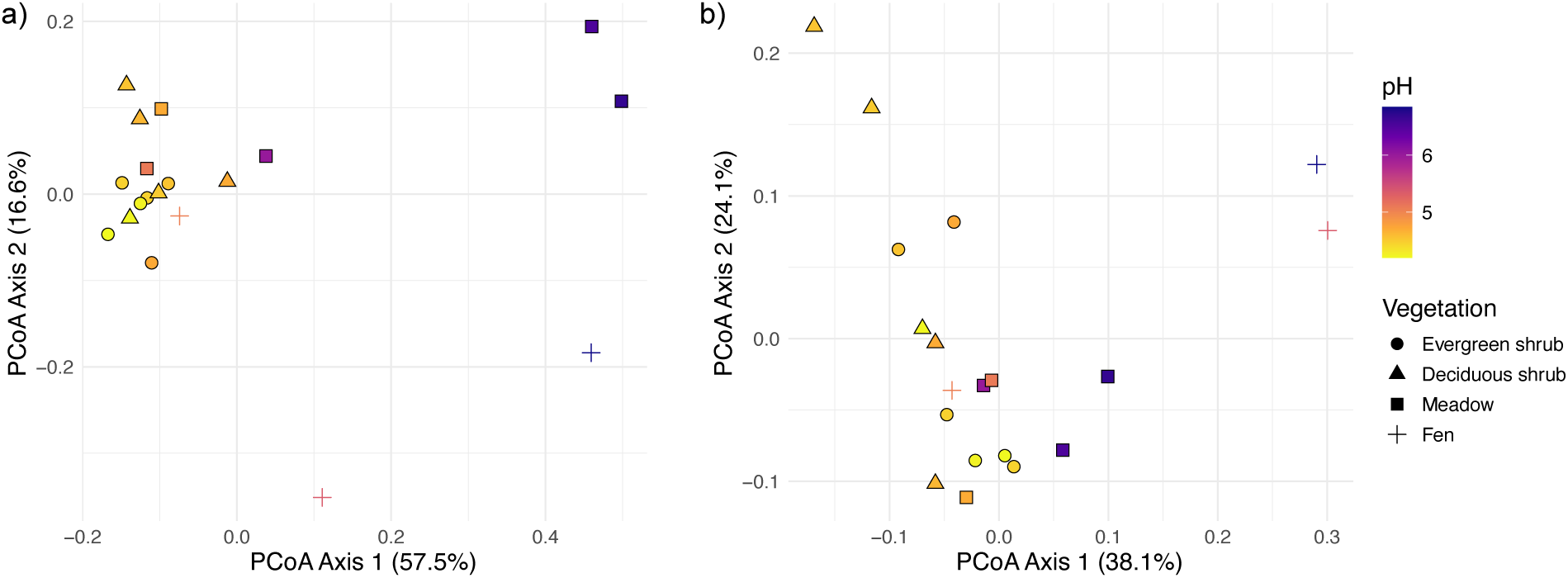
Principal coordinates analysis (PCoA) ordination of active oroarctic soil bacterial and archaeal community composition based on small-subunit rRNA (SSU rRNA) transcripts (a) and transcribed protein-coding genes with KEGG annotation representing the functional composition (b) based on Bray-Curtis dissimilarity. Each point represents a sample, with colors indicating pH and shapes representing vegetation type. The axes (PCoA1 and PCoA2) represent the primary dimensions of variation in the dataset, with the percentages in parentheses indicating the proportion of total variation explained by each axis.

### The transcription of functional genes showed microbial activity in winter soils

#### Basic metabolism and stress response genes were widely transcribed

After filtering the rRNA, 23% of the non-rRNA, representing 1% of the total reads, were mapped to the KEGG database. Based on reads mapping to genes in the manually curated functional gene database and KEGG database, central carbohydrate and energy metabolism genes were transcribed, indicating the microbial activity in winter (Figure 4 and Supplementary Figure S3). These included genes encoding cytochrome c oxidase (*coxA*), F-type ATP synthase (*atpA*), aconitate hydratase (*acnA*), and citrate synthase (*gltA*). The relative transcription of several stress-related genes was high in winter (Supplementary Figure S4). These included genes *groEL*/*groES* and *dnaK*/*dnaJ*/*grpE* encoding chaperone systems, *cspA*, *hsp20*, and *ibpA* encoding temperature shock proteins, *gyrA*/*gyrB* and *rhlE* and *deaD* encoding DNA gyrase and ATP−dependent RNA helicases, respectively. Genes *rpoD* and *rpoE* encoding housekeeping sigma factors and the extracytoplasmic stress response sigma factor, and genes *rpoS* and *rpoH* encoding RNA polymerase sigma factors, which control various stress responses, were transcribed in all vegetation types. The relative transcription of genes *otsA* and *otsB,* involved in the biosynthesis of storage polysaccharide trehalose, was higher in deciduous shrubs than in other vegetation types. Gene *phaZ* encoding poly(3−hydroxybutyrate) depolymerase, involved in depolymerization of storage polysaccharides polyhydroxyalkanoates, was transcribed more in meadows than evergreen shrubs (Kruskal-Wallis test, followed by Dunn’s test with Bonferroni correction; *p* < 0.05).

**Figure 4.**
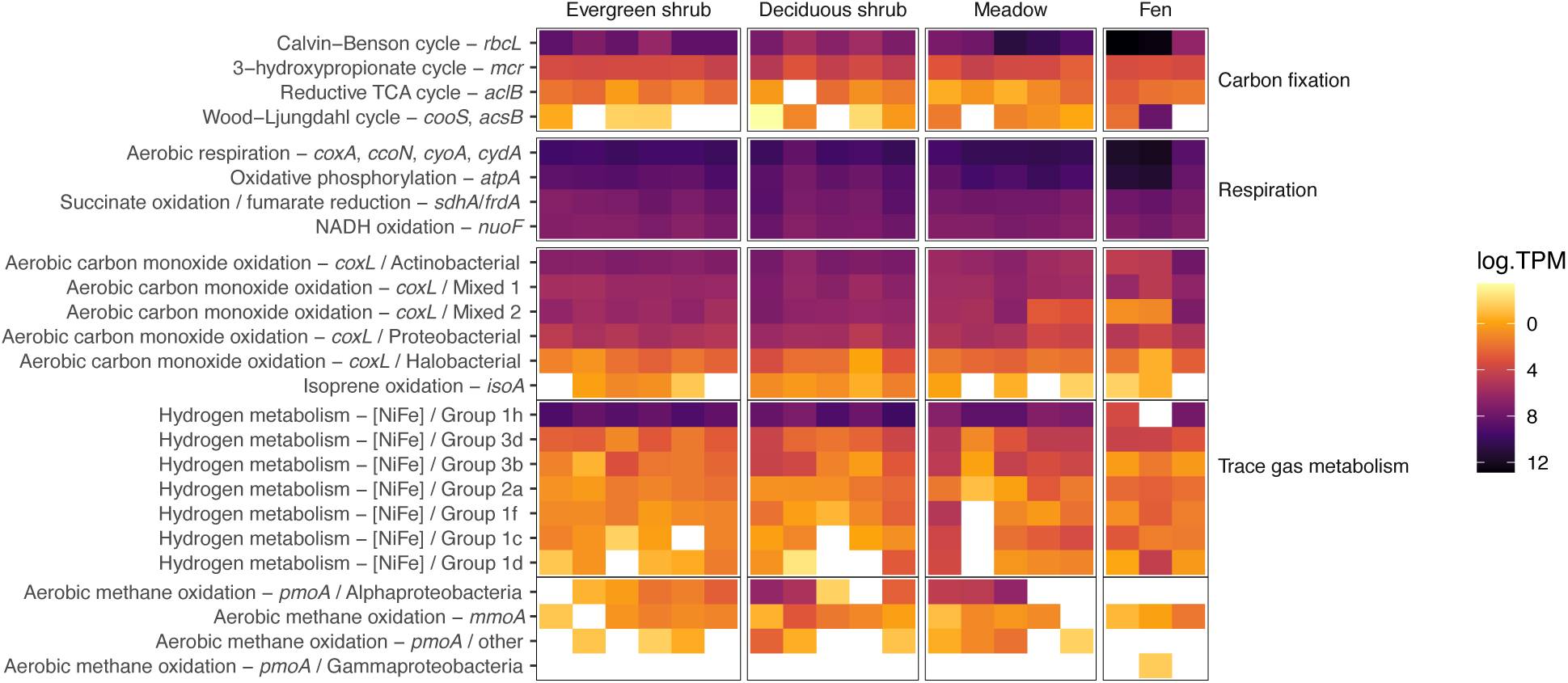
Heatmap showing the relative transcription (log-transformed transcripts per million, TPM) of selected bacterial and archaeal functional marker genes involved in carbon fixation, respiration, and trace gas metabolism in oroarctic tundra soils.

#### Trace gas metabolism may support microbial growth, and N_2_O leaks likely from truncated denitrification during winter

Genes encoding enzymes for the utilization of trace gases methane (CH_4_), carbon monoxide (CO), and dihydrogen (H_2_) were transcribed. The *pmoA* gene encoding particulate methane monooxygenase for CH_4_ oxidation was transcribed in shrubs and meadows (Figures 4 and 5). Based on the curated functional marker gene database, the reads mapped to alphaproteobacterial *pmoA* genes from mainly undefined *Beijerinckiaceae*. This was supported by 16S rRNA data showing alphaproteobacterial methanotroph activity (Supplementary Figure 5). The *mmoA* gene encoding soluble methane monooxygenase was transcribed in all vegetation types. The methanogenesis gene *mcrA* encoding the methyl-coenzyme M reductase was not detected. One fen plot showed high methane emissions, but its metatranscriptomes were compromised. Genes encoding the nickel-iron hydrogenases [NiFe], involved in hydrogen metabolism, were transcribed in all vegetation types (Figure 4). In particular, Actinobacteria-type group 1h [NiFe] was highly transcribed in shrubs and meadows. Groups 3b, 3d, 2a, 1f, 1c, and 1d were transcribed in all soils. Gene *coxL, encoding form I CO dehydrogenase and marking aerobic CO oxidation, was widely transcribed*.

**Figure 5.**
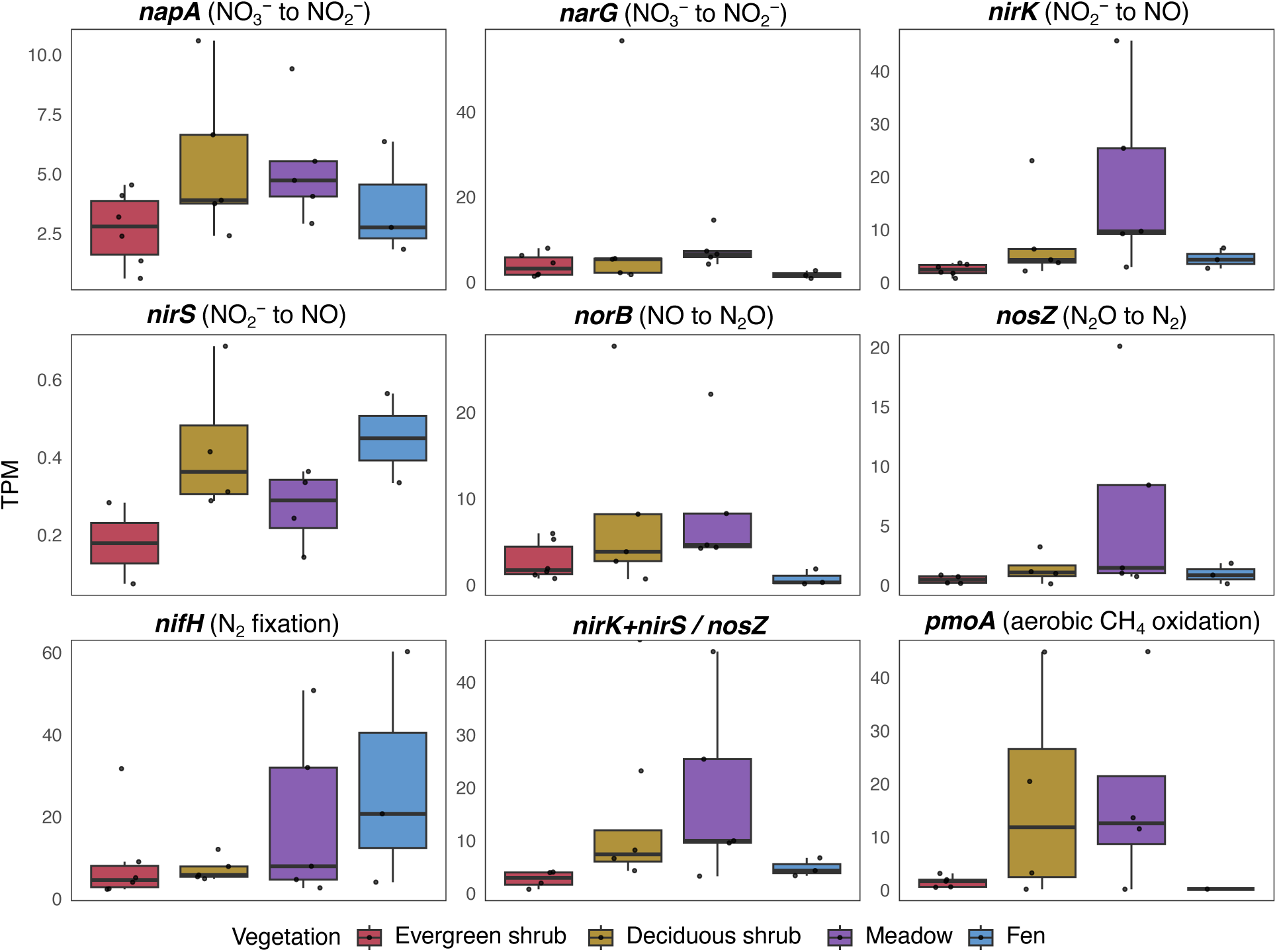
Boxplot showing the relative transcription of functional marker genes involved in denitrification, nitrogen fixation, and aerobic methane oxidation as TPM (transcripts per million) in each vegetation type in the oroarctic tundra soils.

Denitrification genes were transcribed in all vegetation types (Figure 5). The relative transcription of gene *nirK* encoding nitrate reductase was higher in meadows than in other vegetation types (Kruskal-Wallis test, followed by Dunn’s test with Bonferroni correction; *p* < 0.05). The ratio of transcripts for nitrite reductase genes *nirK* and *nirS* to the nitrous oxide reductase gene *nosZ* was not significantly different between vegetation types. However, particular deciduous shrub and meadow samples had higher ratios.

### Statistical models connect gene expression with environmental drivers and GHG fluxes

#### Environmental variables correlated with gene transcription patterns

The activity of most genes was well explained by C/N, GWC, and temperature (Figure 6). The lowest model performance was observed for [NiFe] hydrogenase type 1h, *coxL*, and *mmoA*, which exhibit low TPM variability. The C/N ratio was often the strongest predictor, generally showing a negative effect on TPM. Temperature was the least important variable and often had a negative effect on TPM, suggesting nutrient and moisture availability are stronger controls on gene expression in cold conditions.

**Figure 6.**
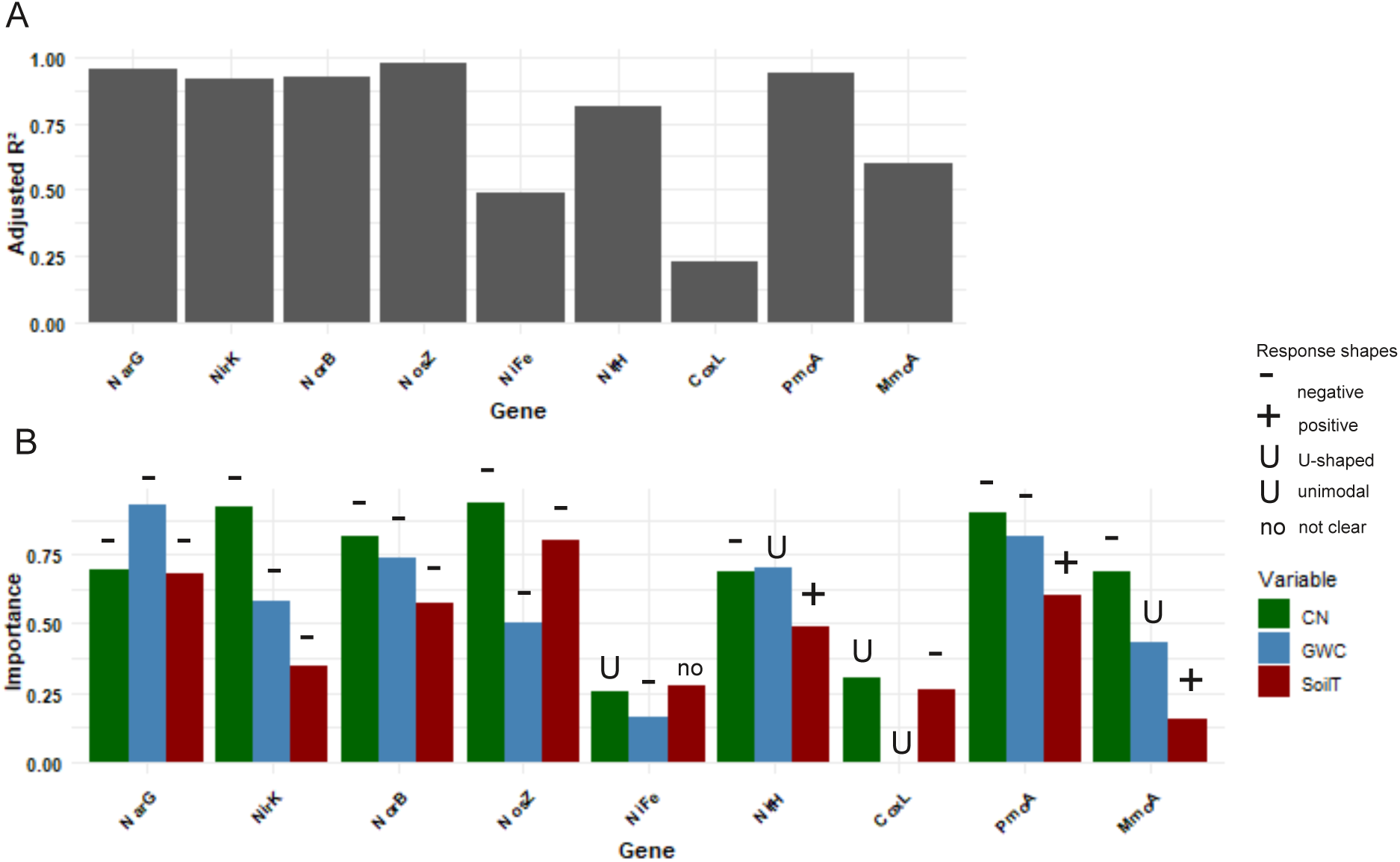
(A) Adjusted R^2^ of the GAM models with TPM as a response variable and GWC, C/N, and temperature as explanatory variables, fit for each gene separately and (B) variable importance scores (bars; 0=low importance, 1=high importance) and the direction of responses (signs above bars) between each response and explanatory variable. An R^2^ value of one indicates a perfect model fit, and a value of zero indicates that the model is completely random. If no bar is shown for the graph, its R^2^ or importance is close to zero.

#### Microbial activity data improve the model performance of wintertime greenhouse gas flux data

Incorporating functional gene data into environmental models explaining GHG fluxes significantly improved model performance, particularly for N_2_O and CO_2_ fluxes. Adding functional gene data increased adjusted R^2^ values from 0.0-0.2 to as high as 0.6 (Fig. 7). For CH_4_ fluxes, R^2^ values were on average 1.4 times higher when functional gene data were included, whereas the increase was 10 and 4-fold for N_2_O and CO_2_ fluxes, respectively. *nirK* and *nosZ* had the greatest impact, while *coxL*, *pmoA*, and *narG* contributed moderately. It should be noted that the relationship between the selected genes and CO_2_ aims to explain only general activity. To explain CO_2_ fluxes more precisely, various C metabolism genes, such as *rbcL,* should be included to draw broader conclusions about CO_2_ production in soils. Nitrogen cycle genes generally showed a positive relationship with N_2_O fluxes, and *pmoA* was positively associated with CH_4_ flux, though weakly.

**Figure 7.**
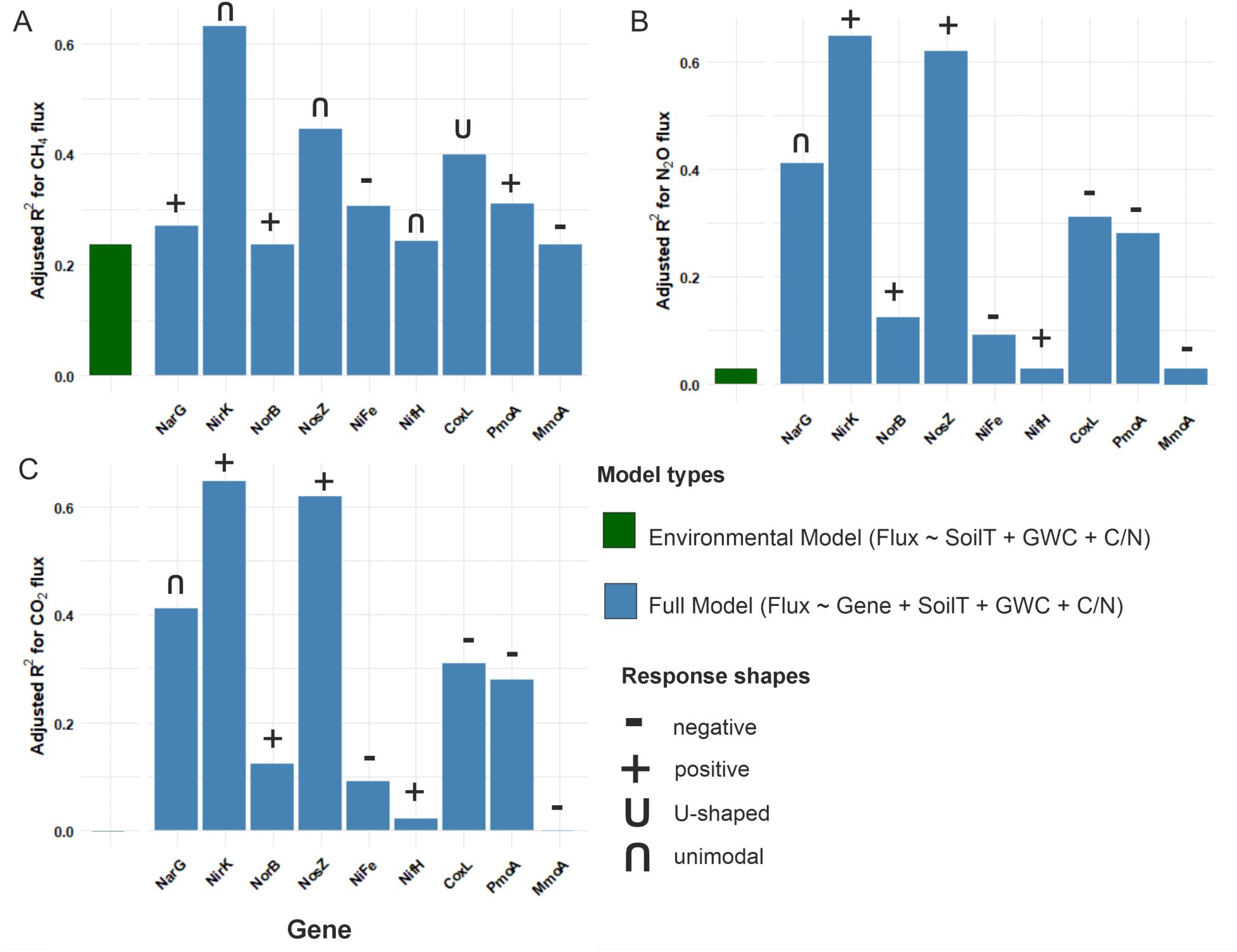
Adjusted R^2^ values for GAM models with GHG fluxes as response variables and TPMs of each gene as explanatory variables: (A) CH4, (B) N2O, and (C) CO2. Green bars represent models using only environmental variables (temperature, GWC, and C/N) as predictors. Blue bars show models that include both environmental variables and the expression of individual functional genes as predictors. The signs above bars show the response direction between each flux and gene.

## DISCUSSION

### Microbial communities are active and contribute to GHG emissions in winter

The transcription of metabolism and stress-related genes, along with detected CO_2,_ N_2_O, and CH_4_ fluxes, indicated microbial activity during winter. CO_2_ fluxes generally fell within the range of winter emissions reported for tundra ecosystems (Treat, Bloom and Marushchak 2018; Natali *et al*. 2019). No significant differences were found between plots; however, deciduous shrubs tended to have higher CO_2_ emissions compared to other vegetation types. Higher CO_2_ emissions may be explained by the substrate quality and quantity: litter from deciduous shrubs is typically more readily degradable than that from evergreen shrubs, as it contains less recalcitrant lignin and fewer microbe-inhibiting tannins and phenolic compounds (e.g., McLaren *et al*. 2017). Meadows with high graminoid and forb cover generally have more easily degradable plant litter (Hobbie 1996), and based on previous studies (Björkman *et al*. 2010; Morgner *et al*. 2010), lower CO_2_ emissions compared to shrubs were not expected. Other factors related to winter conditions, such as the availability of liquid water, soil structure, and reduced priming due to low plant activity, may influence microbial communities, leading to less effective decomposition in meadows.

All soils emitted small amounts of N_2_O, typical for wintertime (Sommerfeld, Mosier and Musselman 1993; Alm *et al*. 1999). Given the long non-growing season in northern tundra regions, they may contribute substantially to the annual N_2_O budget (Rautakoski *et al*. 2024). In fens, high winter water content may lead to nutrient accumulation, providing substrate for denitrification. Our results indicate that nitrogen primarily undergoes denitrification, but some N_2_O escapes into the atmosphere. Despite improved understanding of N_2_O production, controlling factors remain unclear. Emissions of N_2_O from soils are complex (Butterbach-Bahl *et al*. 2013), and the role of dry tundra soils in global N_2_O emissions under changing conditions remains unclear. Since moisture affects N_2_O cycling, examining N_2_O emissions throughout the year could provide insights into the impact of environmental variables on these sporadic N_2_O fluxes and how they are cycled during the changing shoulder seasons.

### Vegetation types drive the microbial community composition

As hypothesized, the active bacterial and archaeal community composition varied between the vegetation types, consistent with Viitamäki *et al*. (2022). Differences were observed along the pH, C/N ratio, and moisture gradients from evergreen shrubs to deciduous shrubs, meadows, and fens. C/N ratios correlated with pH and SOM, reflecting shifts in organic matter quality and soil chemistry that influence microbial niches across vegetation types. The communities were most heterogeneous in meadows and fens, with the highest pH resulting from water flow that accumulates nutrients in these ecosystems.

Four active phyla, *Actinobacteriota*, *Pseudomonadota*, *Acidobacteriota*, and *Planctomycetota*, dominated soils during winter. Orders *Acidothermales*, *Rhizobiales*, and *Acidobacteriales* were highly active in shrub tundra and decreased along the pH gradient. These tundra-soil-dominating heterotrophs (Männistö, Tiirola and Häggblom 2007; Viitamäki *et al*. 2022) likely play a key role in the decomposition of organic matter during winter.

### Trace gas metabolism complements microbial energy demands

Negative CH_4_ fluxes and transcription of aerobic methane oxidation genes indicated small CH_4_ uptake in shrubs and meadows. In fens, methane emissions were negligible, excluding one fen plot with high methane emissions. Although no methanotrophic or methanogenic activity was observed in fens, methane may be produced and consumed in deeper soil layers, even during winter. Methane can be trapped under frozen soil and released in sudden bursts, particularly during autumn (Mastepanov *et al*. 2008; Pirk *et al*. 2017). If aerobic methane oxidation continues throughout the cold season in tundra soils, some trapped CH_4_ is prevented from releasing during spring thaw. Nevertheless, CH_4_ oxidizers may be dormant in certain winter conditions. This kind of spatial and temporal variability is difficult to observe without studying the soil microbial communities, snowpack and soil freezing dynamics, and gas fluxes throughout the seasons.

Active aerobic methanotrophs in shrubs and meadows were mostly *Alphaproteobacteria*, including acidophilic genera *Methylocella*, *Methylocystis,* and *Methylosinus*, which are commonly found in acidic soils. These organisms were also active in less acidic fens despite the absence of detectable methane uptake or oxidation genes. Generally, aerobic methanotrophs oxidize methane as their primary or sole carbon and energy source. Known for their high-affinity methane oxidation, they thrive in upland soils at low CH₄ concentrations (Knief, Lipski and Dunfield 2003; Knief and Dunfield 2005; Tveit *et al*. 2019). Nevertheless, recent studies demonstrate their metabolic versatility: many can utilize multiple carbon sources, such as methanol and acetate, and fix dinitrogen (Dedysh, Knief and Dunfield 2005; Tikhonova *et al*. 2021), and even conserve energy from atmospheric H_2_ and CO (Tveit *et al*. 2019, 2021; Hakobyan *et al*. 2020; Schmitz *et al*. 2020; Schmider *et al*. 2024).

Previous studies have shown that dry upland soils are small CH_4_ sinks during the growing season, for example, in this same area in northern Finland (Virkkala *et al*. 2024), in Greenland (D’Imperio *et al*. 2023) and in Canada (Voigt *et al*. 2023). In contrast, dry Arctic tundra was found to be a notable CH_4_ source during the cold season (Zona *et al*. 2016; Treat, Bloom and Marushchak 2018). In the study, the highest CH_4_ fluxes were observed in the fall, whereas fluxes were close to zero during January-May. Our results on the transcription of methane oxidation genes and gas measurements suggest that oroarctic tundra soils may even act as a small sink during the time of deepest snow cover.

Genes encoding [NiFe] group 1h hydrogenases were widely transcribed in Kilpisjärvi soils during winter, suggesting microbes utilized atmospheric H_2_ for energy during carbon scarcity. This hydrogenotrophic respiration is common in oligotrophic soils, particularly in aerated soils that contain trace concentrations of H_2_. It is found in aerobes and facultative anaerobes across the phyla *Acidobacteriota*, *Actinomycetota, Verrucomicrobiota*, *Chloroflexota*, and *Planctomycetota* (Greening *et al.,* 2014, 2015a, 2015b; Giguere *et al.,* 2020; Bay *et al.,* 2021). Recently, a novel potential nitrogen-fixing *Eremiobacterota* MAG with the potential for H_2_ utilization was characterized in a study from the same plots (Pessi *et al*. 2024). Hydrogenases are also present in methanotrophs, such as *Methylocystis* (Hakobyan *et al*. 2020), *Methylocapsa* (Tveit *et al*. 2019), and *Methylacidiphilum* (Schmitz *et al*. 2020), which may indicate that H_2_ supports their growth. Overall, we showed that H_2_ is a potential energy source for microbes in shrubs and meadows in winter.

Gene *coxL*, encoding the large subunit of form I CO dehydrogenases, was also transcribed widely. As our reads mapped widely to *coxL* from *Actinomycetota*, CO oxidation may be a significant means of survival for this abundant group during winter. Aerobic CO oxidation supports microbial survival during carbon starvation in soils (Cordero *et al*. 2019). Additionally, CO plays a role in atmospheric chemistry, as it reacts with OH radicals, which could otherwise reduce the amount of other GHGs. Along with this chemical consumption, microbial oxidation of CO is a major sink, particularly in surface soils (King 1999). Soil and air temperature, soil moisture, and SOC are the primary controllers of CO oxidation globally, and increasing SOC increases the CO substrate (Liu *et al*. 2018). Bay *et al*. (2021) found that H_2_ and CO are rapidly oxidized in diverse soil ecosystems, while methane is oxidized to a lesser extent, which may also be true in oroarctic soils.

### Microbial activity data are needed to understand wintertime GHG flux dynamics

Environmental variables explained transcription well, demonstrating that wintertime microbial activity can be effectively modeled. Moreover, incorporating microbial data was crucial for explaining winter GHG fluxes, as environmental variables alone often provided limited explanatory power. Notably, nitrogen cycle genes such as *nirK* and *nosZ* strongly influenced N_2_O fluxes, while *pmoA* played a key role in CH_4_ uptake. These findings underscore the importance of integrating metatranscriptomic data into biogeochemical and ecosystem models — a technique still in its infancy, not only in winter studies. Although promising studies have begun to explore the functional implications of microbial communities for GHG fluxes (e.g., Oh *et al*. 2020), major gaps remain in our ability to predict and model microbially mediated processes in tundra soils (Schädel *et al*. 2024). Our results are a promising step in this direction, demonstrating that while environmental data can effectively explain wintertime microbial activity, metatranscriptome data are essential for explaining GHG fluxes.

## ACKNOWLEDGEMENTS

SV was funded by the University of Helsinki’s Doctoral Program in Microbiology and Biotechnology, and JH was funded by the Research Council of Finland (project DARKFUNCTIONS, Grants no. 314114 and 335354 and project N-FUNK, Grant no. 354462). Christina Biasi was supported by the Austrian Science Fund (project PERNO, no. 10.55776/M3335) and the Research Council of Finland through multiple projects, including N-PERM (no. 341348), NOCA (no. 314630), and the Yedoma-N (no. 287469). MEM received funding from the Research Council of Finland (Thaw-N; Grant no. 353858). AMV acknowledges funding catalyzed by the TED Audacious Project (Permafrost Pathways). Metsähallitus granted permission to perform fieldwork. We acknowledge the Kilpisjärvi Biological Station and its staff for the opportunity to use their premises. During the preparation of this manuscript, we used AI tools to assist with editing and analysis. Grammarly was used to enhance grammar and clarity, and ChatGPT (OpenAI) and Microsoft Copilot were utilized to support the editing of written content and the development of R code for data analysis. All outputs generated by these tools were reviewed and verified by the authors.

## Notes

### Competing Interest Statement

The authors have declared no competing interest.

