## Supplementary material for "Microbial community composition explains wintertime greenhouse gas fluxes in an oroarctic tundra ecosystem"

Supporting Information

Figures S1, S2, S3, S4, S5

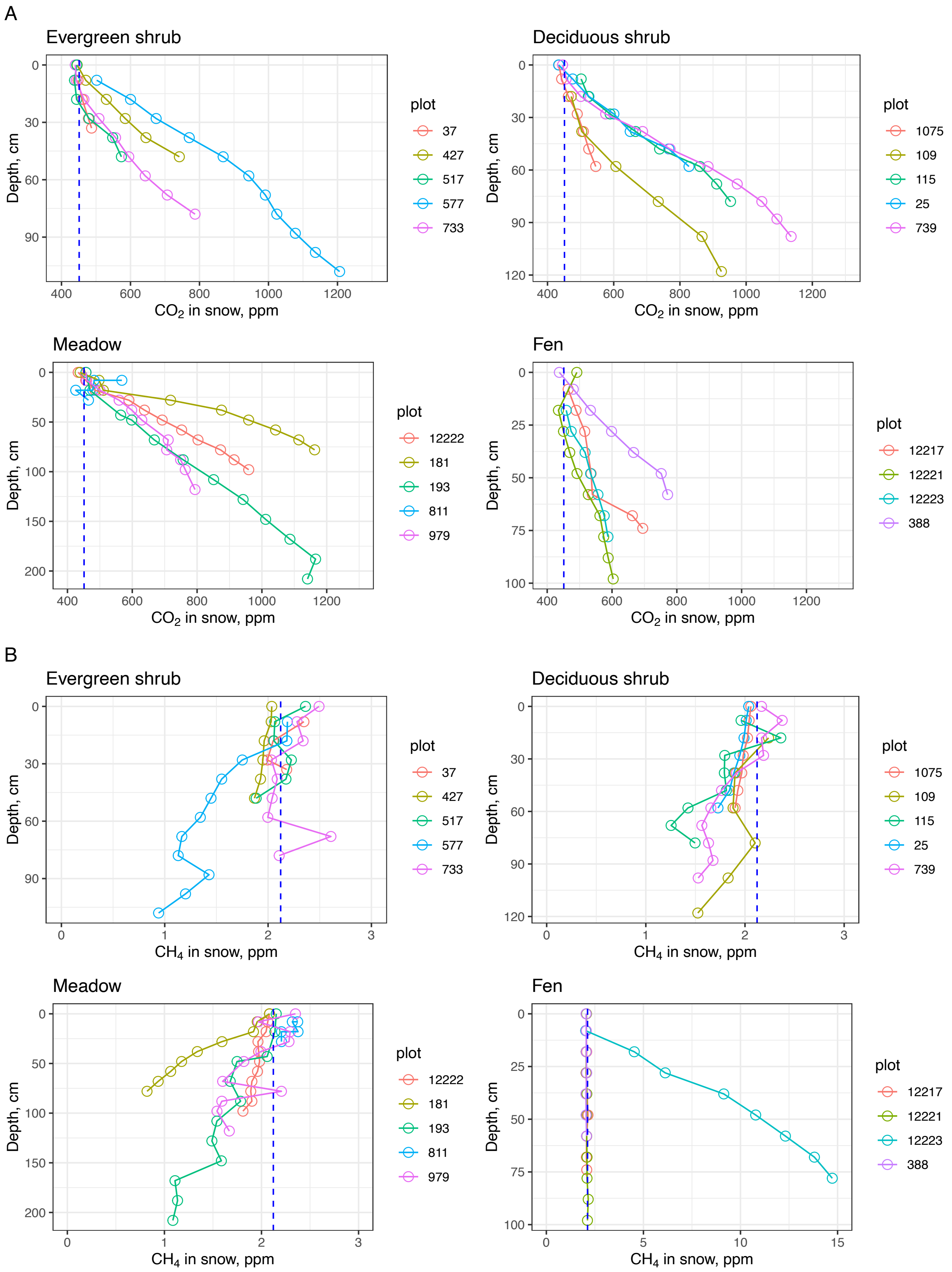

A
Evergreen shrub
Deciduous shrub
Meadow
Fen
Depth, cm
$CO_2$ in snow, ppm
plot
37
427
517
577
733
1075
109
115
25
739
12222
181
193
811
979
12217
12221
12223
388
B
Evergreen shrub
Deciduous shrub
Meadow
Fen
Depth, cm
$CH_4$ in snow, ppm
plot
37
427
517
577
733
1075
109
115
25
739
12222
181
193
811
979
12217
12221
12223
388

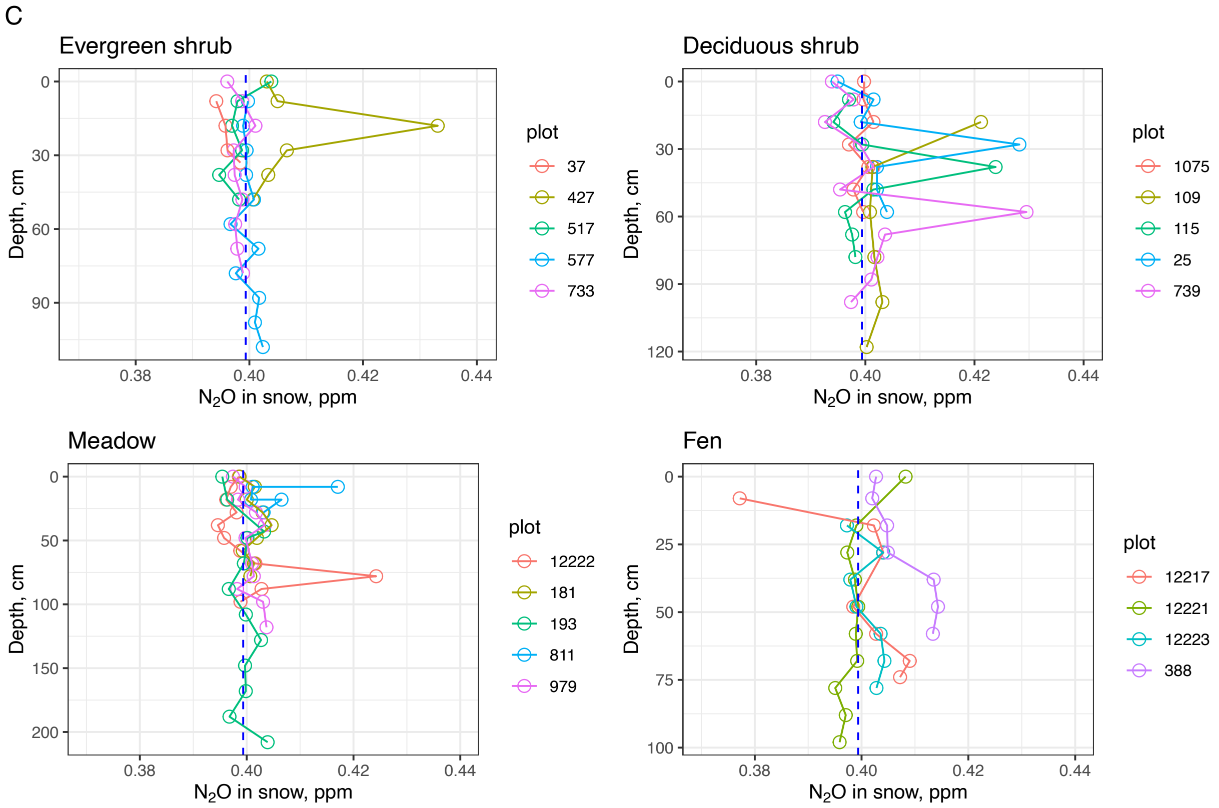


*Supplementary Figure S1. GHG gas profiles of A) $CO_2$, B) $CH_4$, and C) $N_2O$ across the sampled depths in each vegetation type. The ambient gas concentration is represented by a dashed blue line, indicating when concentrations exceed the ambient (net emission) and when they fall below the ambient (net consumption).*

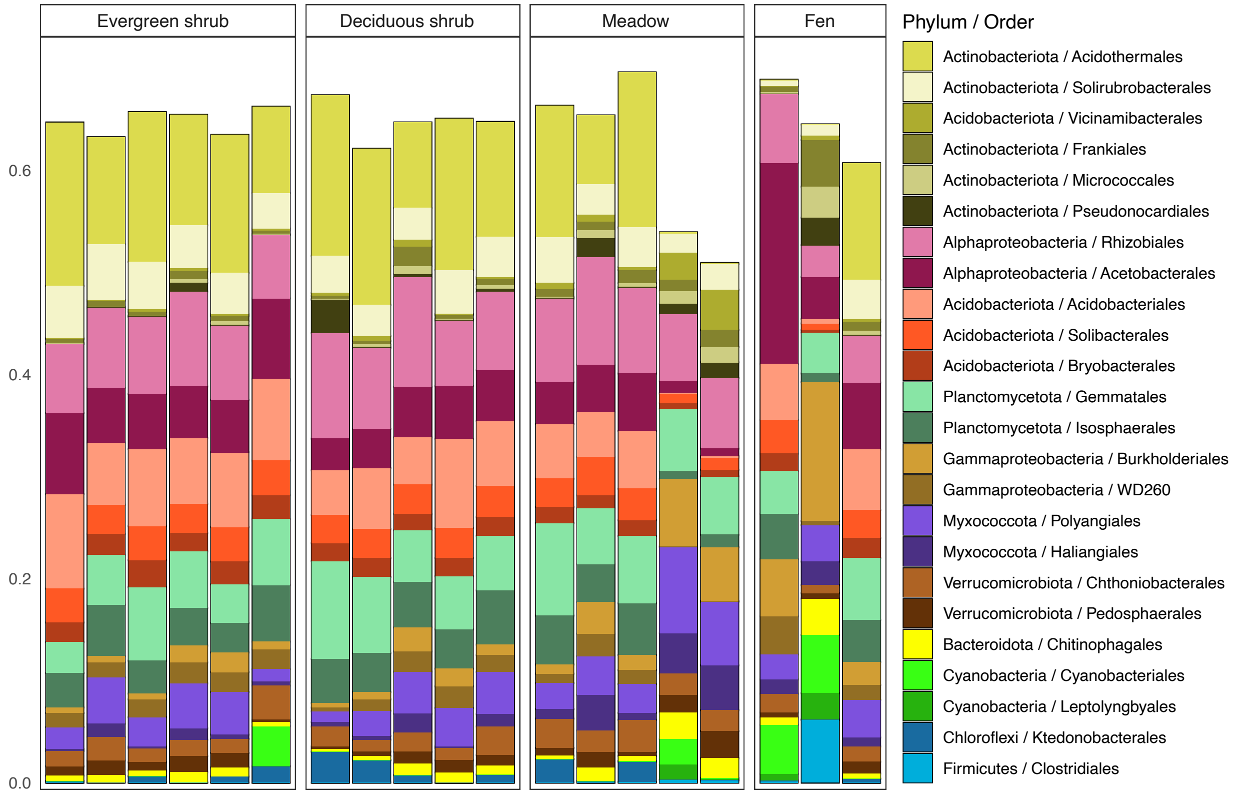


*Supplementary Figure S2. Barplot showing the cumulative relative abundance of bacterial and archaeal taxa at the order level in each vegetation type. Note that* Alphaproteobacteria *and* Gammaproteobacteria *are included at the class level, although they are currently classified under the phylum* Pseudomonadota*. Active taxa representing over 0.025% in a sample are shown.*

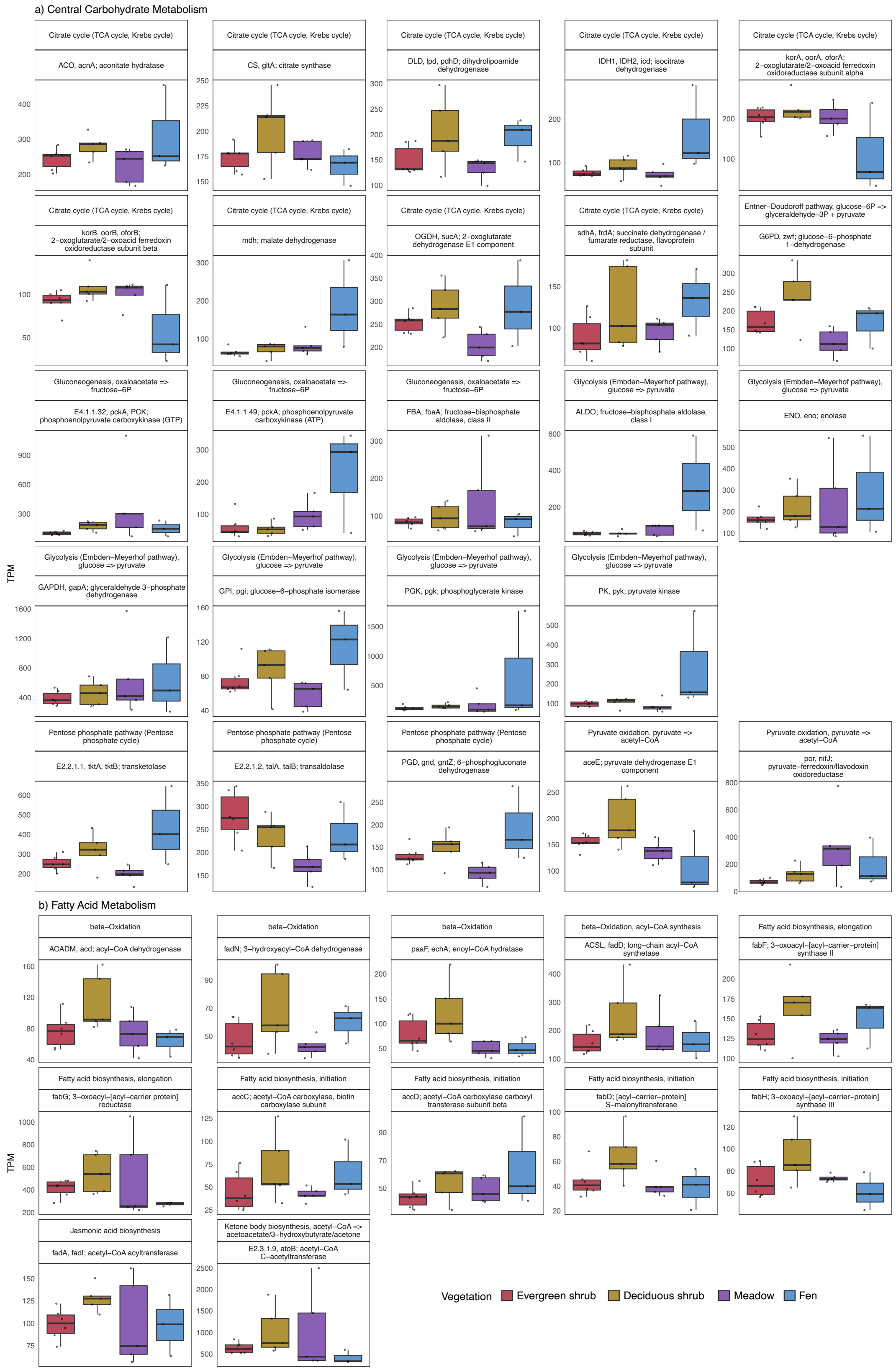


*Supplementary Figure S3. Boxplots showing the most transcribed central metabolism genes (a) and fatty acid metabolism genes (b) in each vegetation type as transcripts per million (TPM).*

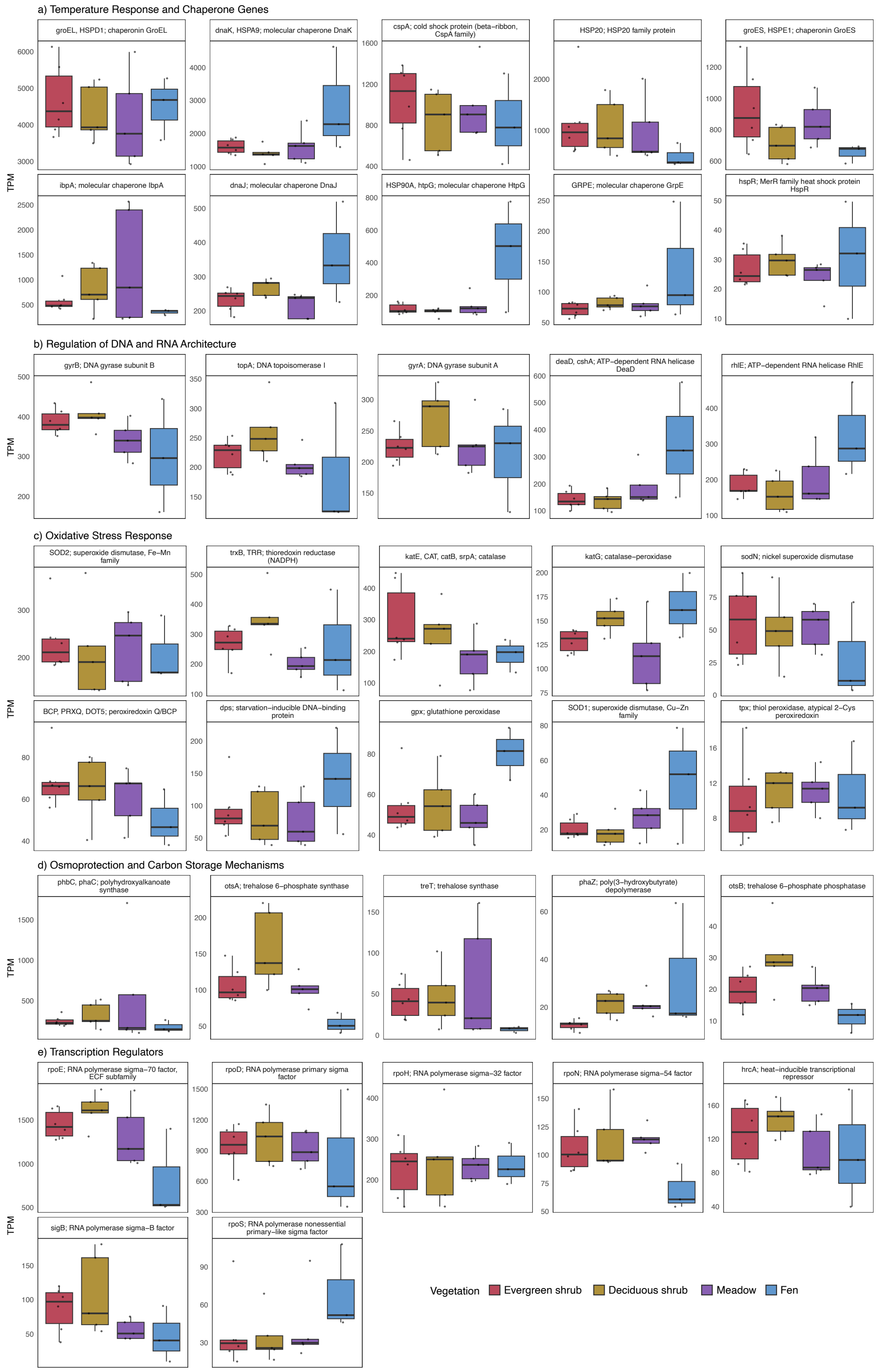


*Supplementary Figure S4. Boxplots showing the selected transcribed temperature response and chaperone genes (a), genes involved in the regulation of DNA and RNA architecture (b), oxidative stress response genes (c), genes involved in osmoprotection and carbon storage mechanisms (d), and transcription regulation genes (e) in each vegetation type as transcripts per million (TPM).*

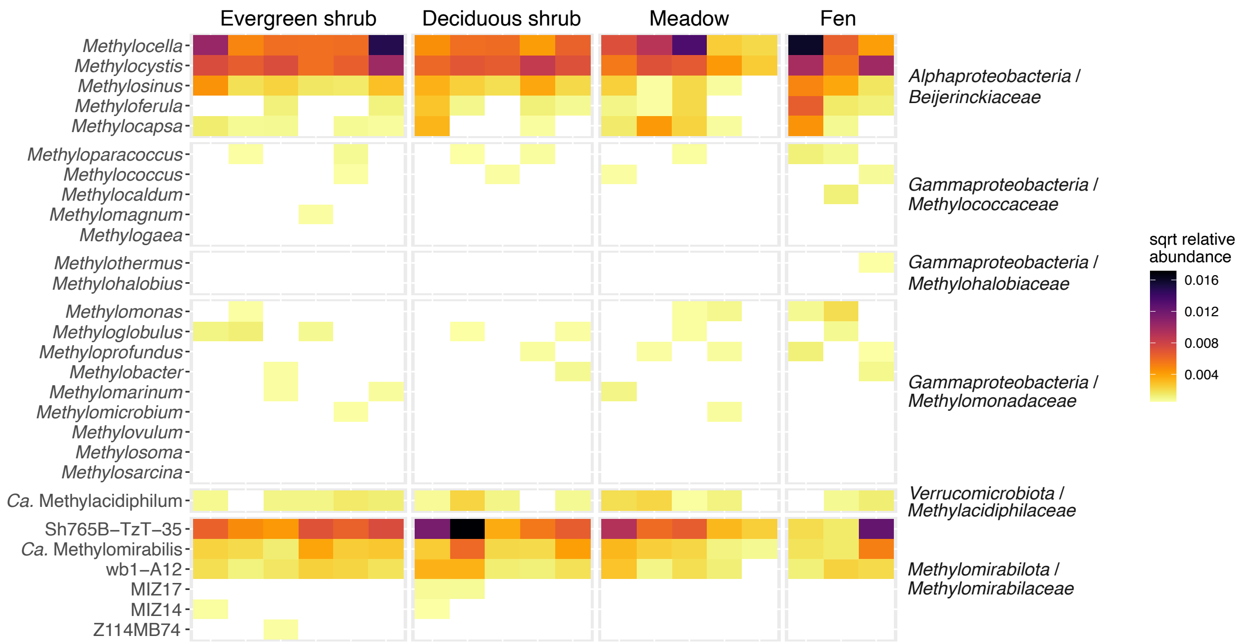


*Supplementary Figure S5. Heatmap showing the square-root transformed relative abundance of the 16S rRNA gene of methanotrophic* Alphaproteobacteria*,* Gammaproteobacteria*, and* Verrucomicrobiota*, and potentially methanotrophic* Methylomirabilota*.*
